# New Insights on Bridging Integrator 1 Protein Isoforms as a Risk Increasing Gene in Alzheimer’s Disease

**DOI:** 10.1101/2025.03.31.646439

**Authors:** Min Hu, Sahar Esmaeeli, Lindsay Stolzenburg, Mariam Jouni, Eugene Nyamugenda, Peter Reinhardt, Lamiaa Bahnassawy, Laura Gasparini, Jeffrey F. Waring, Aparna Vasanthakumar

## Abstract

**INTRODUCTION:** Genome-wide association studies (GWAS) have identified Bridging Integrator 1 (*BIN1*) as the second-most significant genetic risk factor for Alzheimer’s disease (AD). We performed GWAS by proxy (GWAX) using UKBiobank and replicated this finding. However, the mechanism by which *BIN1* impacts AD risk is largely unknown.

**METHODS:** To address this, we first measured the expression of *BIN1* isoforms in a human induced pluripotent stem cells (hiPSC) model and further evaluated whether the *BIN1* risk loci associated with the expression of *BIN1* isoforms in a Phase 2 AD clinical trial.

**RESULTS:** Our data indicated *BIN1* isoform expression patterns associated with the differentiation of hiPSC into neurons or microglia and found the variant rs35103166 impacts the expression of the *BIN1* microglia-specific isoforms.

**DISCUSSION:** Given the strong association with susceptibility to AD, exploring the mechanisms of *BIN1* genetic variants and their impacts on cell-type specific expression could serve as a valuable resource for novel drug discovery.

**Highlights:**

- *BIN1* variants are significantly associated with AD risk in GWAX analysis from UKBB.
- *BIN1* isoforms show a cell-type-specific expression pattern during hiPSC differentiation into neurons or microglia.
- *BIN1* risk alleles associate with *BIN1* isoform expression in a Phase 2 AD trial (NCT02880956).

## 1 BACKGROUND

Alzheimer’s disease (AD) is a complex neurodegenerative disorder associated with age characterized by progressive cognitive decline, memory loss, and impaired daily functioning [1,2]. Pathologically, intracellular neurofibrillary tangles of hyperphosphorylated tau and extracellular β-amyloid plaque deposition are two hallmarks in AD brain [3–7]. AD is the most common cause of dementia and accounts for up to 80% of all dementia diagnoses [8,9], affecting millions of individuals worldwide and increasing healthcare cost dramatically [10–12]. Despite extensive research efforts, there is significant unmet medical need for AD treatment. Therefore, there is an urgent need to gain a deeper understanding of the molecular mechanisms involved in AD pathogenesis and disease progression to identify novel therapeutic targets for treating AD.

Studies show that between 60–80% of the risk of developing AD is driven by heritable factors [9,13]. Several large genome-wide association studies (GWAS) have identified more than 80 AD-associated genetic risk loci [14–18]. The Bridging Integrator 1 Protein (*BIN1*) was found to be the second-most significant genetic risk factor for AD after apolipoprotein E (*APOE*) [14,16,18–23].

The *BIN1* gene is located on chromosome 2q14.3. It has 20 exons and 14 isoforms derived from alternative splicing of its pre-mRNA, and 11 of these 14 RNA transcripts translate into proteins [24–26]. These isoforms exhibit distinct cellular localizations [24,27–29]. Although not all research findings indicate consistent isoform distribution patterns, literature indicates that *BIN1* isoforms 1, 2, 3, 5 and 7 are expressed in brain, primarily in neurons and astrocytes, referred to as neuronal-astrocytic-specific isoforms [26,28,30]. *BIN1* isoforms 6, 9, 10 and 12 are predominantly found in microglia, so they are referred to as microglia-specific isoforms [26,31]. *BIN1* isoforms 4 and 8 are not expressed in the central nervous system [26,32], and isoform 8 is mainly expressed in the muscle [30,32]. This cellular distribution pattern suggests that *BIN1* isoforms may have distinct functions in specific cell types and specific brain regions, contributing to the complexity of AD pathogenesis [19,33].

Several studies demonstrate that *BIN1* contributes to AD pathology by either mediating the production and intracellular aggregation of Aβ42 [34–39] or by modulating tau pathology [19,26,40–44]. Some studies observed a decrease of *BIN1* isoform 1 and an increase of isoform 9 expression in human AD post-mortem brain samples [41,42].

Single nucleotide polymorphisms (SNPs) in *BIN1* gene also have been studied for their association with AD risk. Multiple *BIN1* SNPs including rs59335482 [19], rs74490912 [26], rs7561528 [23,45,46], rs6431223 [43]), rs6733839 [18,43], rs754834233 [47] and rs138047593 [47] have been identified and reported to influence AD susceptibility, with rs744373 being the most widely investigated [48–54].

To better understand the precise mechanisms by which *BIN1* genetic variations drive the expression of *BIN1* isoforms and affect AD risk, we first measured the expression of *BIN1* isoforms in human induced pluripotent stem cells (hiPSCs) - derived neurons or microglia. We then characterized how genetic *BIN1* risk alleles influence *BIN1* isoform expression in a Phase 2 AD trial (NCT02880956) and tested their association with AD clinical endpoints in this trial. With this study, we hope to provide new insights on the mechanism by which *BIN1* SNPs and *BIN1* isoforms associate with AD risk, and this may help guide the development of novel therapeutic strategies for AD.

## 2 METHODS

### 2.1 Stratified case-control association mapping of Alzheimer’s disease using family history from UK Biobank (UKBB)

A GWAX study using imputed genotypes on the SAIGE (Scalable and Accurate Implementation of Generalized mixed model) platform was conducted, utilizing the family history of AD from UKBB. We thus defined individuals who are not adopted and reported having at least one parent or sibling with AD as proxy cases, while the proxy super controls included subjects 70 years or older who reported no parental history of AD, depression or Parkinson’s disease. Anyone reporting neurological or behavioral disorders was excluded from the control set. We stratified the cases and controls based on their *APOE* status and performed separate association studies on the subjects who are carriers of *APOE ε4*.

### 2.2 hiPSC-differentiation

We used BIONi010-C-13 derived from fibroblast cells using episomal reprogramming (available through EBiSC: www.ebisc.org). Detailed protocols can be found in supplementary materials and methods.

### 2.2.1 hiPSC-derived neurons

hiPSCs were cultured with daily media change and differentiated into cortical neural progenitor cells (NPC) through a series of media changes for 17 days. NPCs were further differentiated into neurons after plating in neuronal maintenance media with special supplemental reagents. Two days after neuronal plating, the cells were treated with 0.15 µg/mL mitomycin C to kill dividing cells. Cells for RNA analysis were collected at various differentiation stages, including the hiPSC stage, NPC stage, and neuron. Neuronal differentiation from the same hiPSC line were performed independently 4-6 times.

### 2.2.2 hiPSC-derived microglia

Generation of microglial-like cells was based upon the method by Haenseler et al. [55]. Embryoid bodies generated by dissociating hiPSCs into single cells were plated according to manufacturer’s suggested protocol for 15-18 weeks to generate pre-microglia. These pre-microglia were pooled, counted, and then plated and differentiated into microglial-like cells with microglia differentiation media. Within one to two weeks, differentiating microglial-like cells begin to exhibit typical morphological features. Microglia-like cells were collected for RNA extraction in Trizol (Thermo Fisher Scientific, MA, USA) on day 14 of differentiation. Microglial differentiation from the same hiPSC line were performed independently up to 9 times.

### 2.3 Primer design for *BIN1* isoforms

Specific primers and probes for *BIN1* isoforms were designed based on Taga et al’s publication [26] using Primer 3 and synthesized by Integrated DNA Technologies (Coralville, IA, USA). Each primer pair and probe are located at the unique exon-exon boundary for that specific *BIN1* isoform with the exception of primer & probe 10 & 12, which detect both isoform 10 and 12, because there is no unique design that can distinguish between these 2 isoforms. See Supplementary Table S1 for detailed information for each primer & probe.

### 2.4 Droplet digital polymerase chain reaction (ddPCR) assay for measuring expression of *BIN1* isoforms

Total RNA was extracted from hiPSCs, pre-microglia, and microglia using the Direct-zol™ RNA MiniPrep Kits (Zymo Research, CA, USA). RNA extraction for neuronal differentiation cells (hiPSCs, NPC, neurons) was done using the RNeasy Mini Kit (Qiagen, MD, USA). RNA from peripheral blood (PB) in the AD trial was isolated using QIAsymphony PAXgene Blood RNA Kit (Qiagen, MD, USA). cDNAs were synthesized using the iScript cDNA Synthesis Kit (Bio-Rad, CA, USA) using 500 ng RNA. ddPCR was performed following the QX200 Droplet Digital PCR System (Bio-Rad, CA, USA) protocol [56–58]. In brief, droplets were generated using the QX200 AutoDG Droplet Digital PCR System (Bio-Rad, CA, USA). PCR amplification was then conducted on a C1000 Touch Thermal Cycler (Bio-Rad, CA, USA) under the following conditions: 95°C for 10 minutes (enzyme activation), 94°C for 30 seconds (denaturation) followed by 59°C for 1 minute (annealing/extension). Denaturation and annealing/extension steps were repeated for 39 cycles followed by incubation at 98°C for 10 minutes for enzyme deactivation. Finally, the reaction was held at 4°C for 30 minutes then 12°C until droplets were ready for reading. Droplets were read using the QX200 Droplet Reader (Bio-Rad, CA, USA) after PCR amplification. Positive and negative droplets were defined and counted based on the detectable level of fluorescence, reflecting the presence of the target molecule in the droplet. The absolute concentration of *BIN1* isoform expression level was calculated by counting the positive and negative droplets and the counts were fitted to Poisson distribution for statistical analysis in QuantaSoft software (Bio-Rad, CA, USA).

### 2.5 Data analysis

All graphs and data analyses were prepared using GraphPad Prism version 10.3 (GraphPad Software, San Diego, CA, USA). All data were shown as mean ± S.D. For the hiPSC differentiated cell line data, ordinary one-way ANOVA with post-testing using Tukey’s multiple comparisons test was used to evaluate statistical differences in the groups. For the PB data from the AD clinical trial, linear regression analyses were performed using age, sex, race, and *APOE* status as covariates. Wherever applicable, *P* value < 0.05 was considered statistically significant.

## 3 RESULTS

### 3.1 *BIN1* variants are significantly associated with AD risk in GWAX analysis from UK Biobank (UKBB)

A GWAX analysis was performed in the UKBB cohort by utilizing the family history of AD. As shown in Figure 1a, after inclusion and exclusion criteria were met, we had 55,806 cases versus 122,538 controls. The GWAX analysis identified the most well-known loci associated with AD, including *APOE*, *BIN1*, *TREM2* etc. (Figure 1b). Upon stratifying the subjects using *APOE* status as a co-variant, we found that across all stratifications (*APOE*4+, *APOE*4-, *APOE*3/3 subjects), the *BIN1* locus was identified as most significantly associated with AD risk (Figure S1).

**Figure 1.**
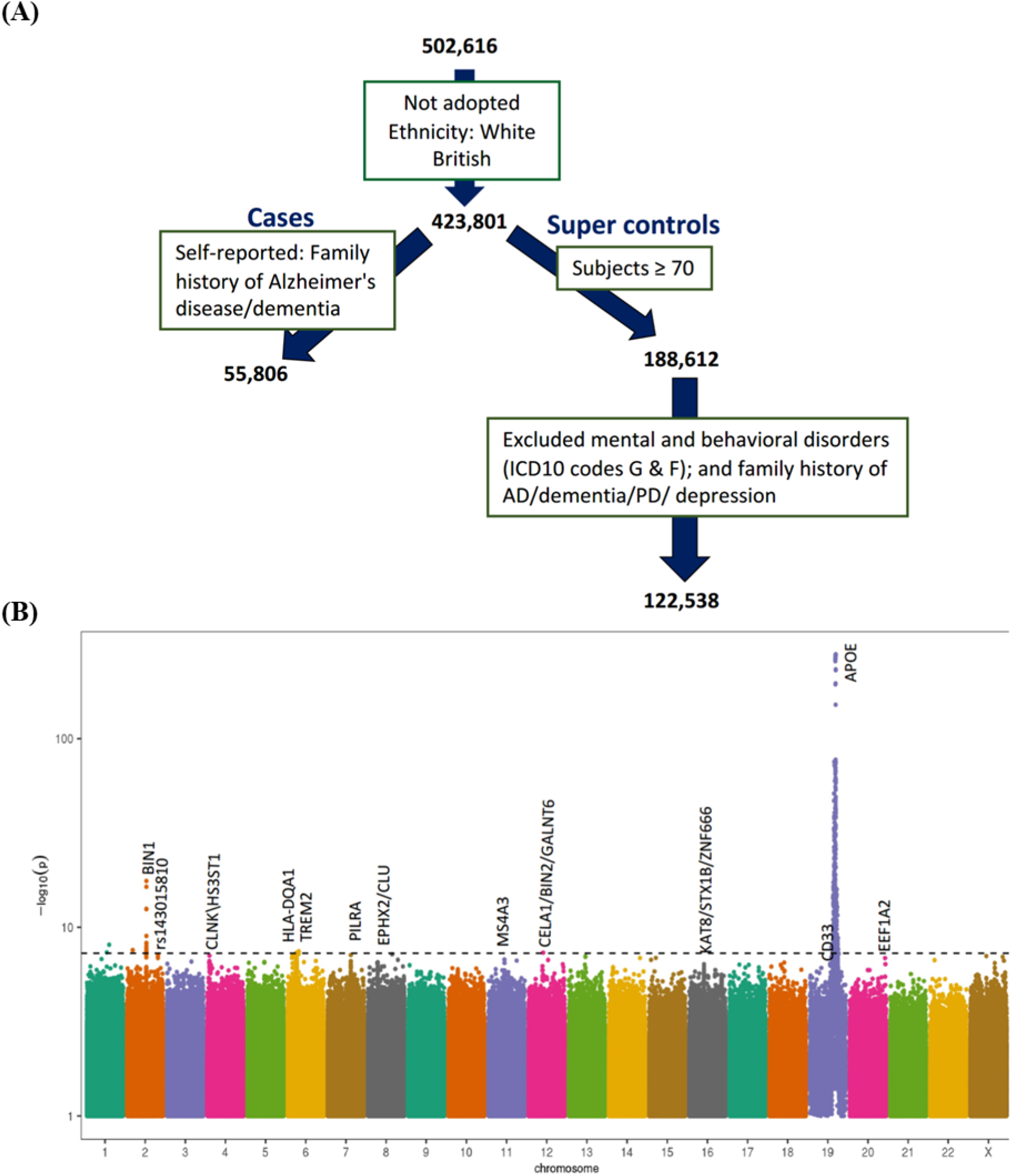
Stratified case-control association mapping of Alzheimer’s disease using family history from UK Biobank (UKBB). (A) A genome-wide association study by proxy (GWAX) using imputed genotypes on SAIGE (Scalable and Accurate Implementation of Generalized mixed model) platform was conducted on 55,806 cases versus 122,538 controls, utilizing the family history of AD from UKBB. Individuals who are not adopted and reported as having at least one parent or sibling with AD were defined as proxy cases, while the proxy super controls include subjects 70 years or older who reported no parental history of AD, depression or Parkinson’s disease. Anyone reporting any neurological or behavioral disorders has been excluded from control set. (B) Manhattan plot showing the genome-wide significant hits from the GWAX analysis with chromosome numbers on the X-axis and genome-wide significance on the Y-axis (log_10_(p-value)), showing APOE (chr. 19) as the top significant hit followed by *BIN1* (chr. 2).

### 3.2 Cell-type-specific expression of *BIN1* isoforms in human iPSC (hiPSC) models

The mechanism by which *BIN1* SNPs alter AD pathology is still unknown. It is hypothesized that *BIN1* variants increase AD risk through their effect on the expression of the *BIN1* isoforms. However, their expression patterns within neuronal cells are not as clear. Therefore, we first investigated the distribution pattern of different *BIN1* isoforms in neuronal and microglial cell types using a hiPSC differentiation model.

During hiPSC differentiation into neurons, we observed that the expression of neuronal-astrocytic-specific *BIN1* isoforms (isoforms 1, 2, 3, 5 and 7) increased significantly, with maximal expression at the neuron stage (Figure 2A and Figure S2A-D). On the other hand, the expression of microglia-specific *BIN1* isoforms (isoforms 6, 9, 10 and 12) increased significantly in the neuronal progenitor cells (NPC), which then decreased to levels similar to hiPSC when differentiated into neurons (see Figure 2B and Figure S2E-F). During the differentiation of hiPSC into microglia, the expression levels of neuronal-astrocytic-specific *BIN1* isoforms decreased during differentiation (Figure 3A and Figure S3A-D). Microglia-specific *BIN1* isoforms significantly increased in microglia relative to both the hiPSC and pre-microglia stages (Figure 3B and Figure S3E-F).

**Figure 2.**
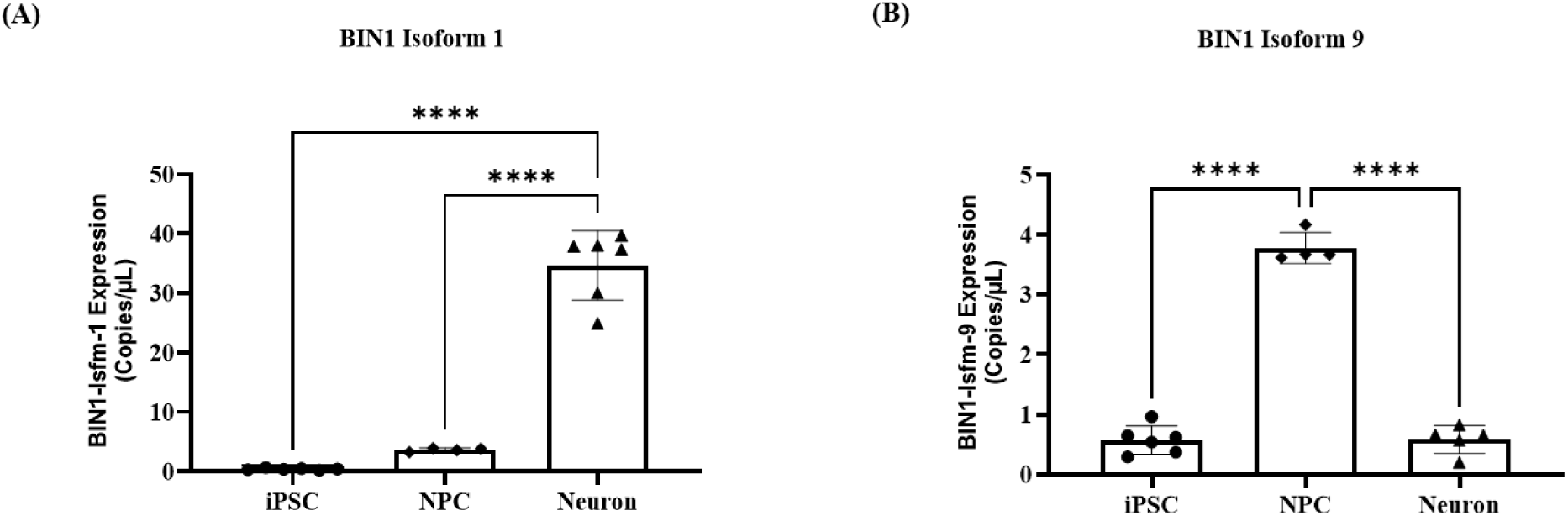
Change in expression level of *BIN1* isoforms during hiPSC differentiation into neurons. hiPSCs were differentiated into neural progenitor cells (NPC), and then into neurons. The expression level of *BIN1* isoforms was measured by ddPCR technology. (A) Expression level of neuronal-astrocytic-specific *BIN1* isoform 1 during differentiation from hiPSCs into NPCs, and finally into neurons. (B) Expression level of microglia-specific *BIN1* isoform 9 during differentiation from hiPSCs into NPCs and finally into neurons. In each panel, X-axis indicates stage of differentiation, Y-axis indicates *BIN1* isoform expression quantified as copies/uL. Each point represents the mean ± S.D. of 4-6 independent experiments. Ordinary one-way ANOVA was performed first to compare between groups with ****, *p*<0.0001. Post-testing was done using Tukey’s multiple comparisons test to compare every group to the other with *, p < 0.05; **, p < 0.01; ***, p < 0.001;****, *p* < 0.001.

**Figure 3.**
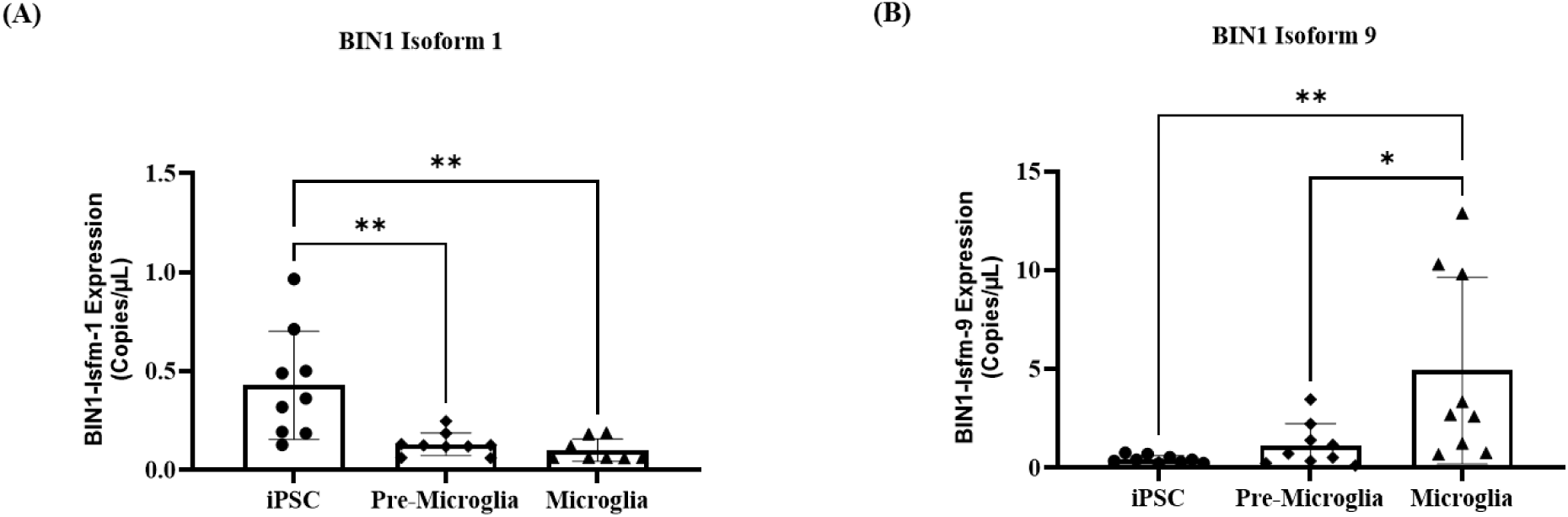
Changes of expression level of *BIN1* isoform during hiPSC differentiated into microglia. hiPSC was differentiated into pre-microglia first, then continued differentiated into microglia. ddPCR technology was applied to detect the expression level of *BIN1* isoform. (A) Expression level of neuron-specific *BIN1* isoform 1 during differentiation from hiPSCs into pre-microglia, and finally into microglia. (B) Expression level of microglia-specific *BIN1* isoform 9 during differentiation from hiPSCs into pre-microglia, and finally into microglia. In each panel, X-axis indicates stage of differentiation, Y-axis indicates *BIN1* isoform expression quantified as copies/uL. Each point represents the mean ± S.D. of 9 independent experiments. Ordinary one-way ANOVA was performed first to compare between groups with ****, *p*<0.0001. Post-testing was done using Tukey’s multiple comparisons test to compare every group to the other with *, p < 0.05; **, p < 0.01; ***, p < 0.001;****, *p* < 0.001.

### 3.3 Association of the *BIN1* risk alleles with *BIN1* isoform expression in a Phase 2 AD trial

To evaluate whether the expression of *BIN1* isoforms associated with the *BIN1* risk SNPs, we utilized samples collected in a clinical trial for early AD (NCT02880956). This was a trial that tested the efficacy and safety of an anti-tau antibody, Tilavonemab, in patients with early AD [59] (Table 1). Genotypes at the *BIN1* loci most significantly associated with risk of AD in our GWAX analyses (rs35103166, rs4663105, rs6733839, rs744373 and rs7561528), all of which are upstream of the *BIN1* gene, were extracted from WGS analyses. Correlation of these SNPs with the expression of *BIN1* isoforms in PB from patients at Day 1 was evaluated. We only detected microglia-specific isoforms in the peripheral blood, given that microglia are brain-specific macrophages. Since our goal was to examine the effect of the alternate allele on downstream expression, we combined the data from the heterozygous and alternate homozygous groups for all our analyses. We found that rs35103166 was significantly associated with increased expression of microglia-specific isoforms (Isoform 6, 9, and 10&12) in patients who harbored the alternate allele (C/C & T/C) compared with patients who had the reference homozygous genotype (T/T) (Figure 4 panel (A), (B) and (C)). The other four *BIN1* SNPs (rs4663105, rs6733839, rs744373 and rs7561528) that we tested, including the most studied rs744373, didn’t show any significant association with *BIN1* isoform expression (Supplementary S4).

**Figure 4.**
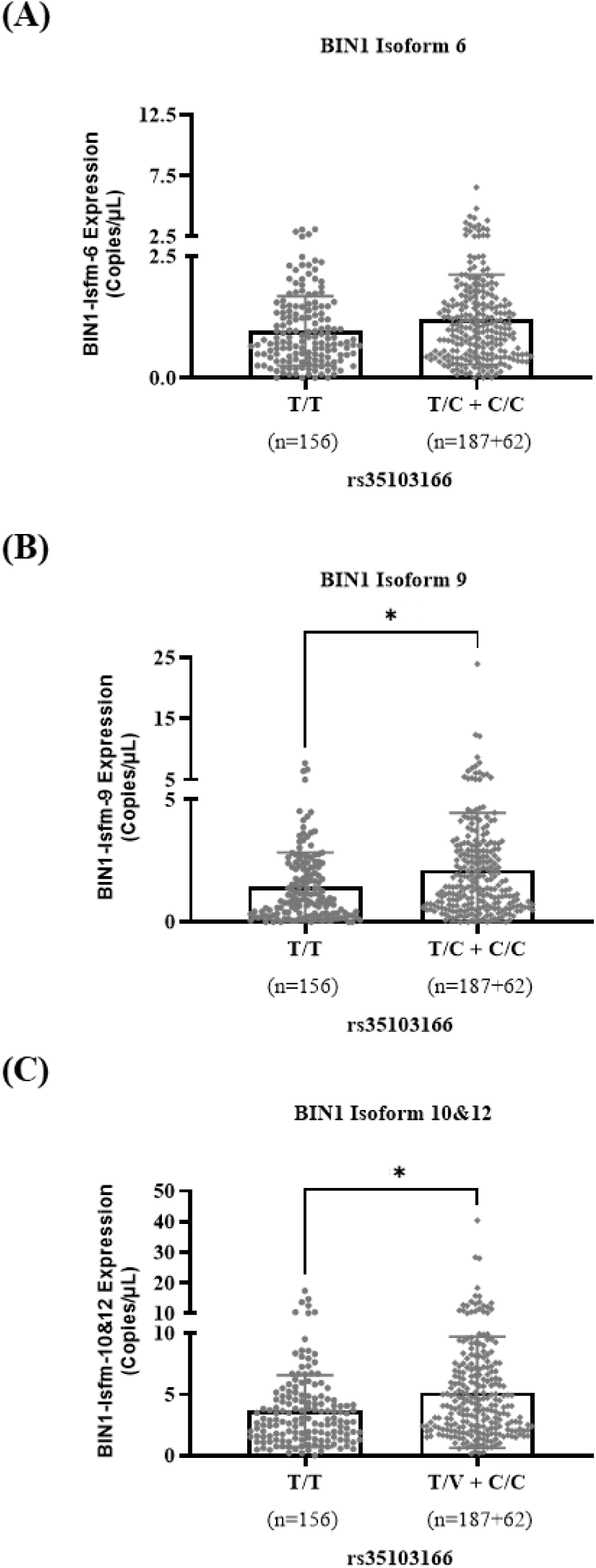
Effect of *BIN1* genotype on expression of *BIN1* isoform in PB from an AD trial. RNA from PB samples collected at baseline from the AD trial (NCT02880956) was used to measure the expression of *BIN1* isoforms using ddPCR technology. The whole genome sequencing data from this trial was used to identify genotype status at *BIN1* SNP rs35103166. At this locus, 156 subjects had reference homozygous (T/T), 187 subjects had heterozygous (T/C) and 62 subjects had alternate homozygous (C/C) genotype. All subjects containing alternate genotypes were grouped together for this analysis, as indicated on the X-axis. Linear regression analysis was performed to test the association of the presence of the SNP with the expression of specific *BIN1* isoforms using age, sex, and *APOE* status as covariates. Each individual panel shows the expression of *BIN1* isoform 6 (A), isoform 9 (B), and isoform 10&12 (C). *p* values were 0.13, 0.04 and 0.01 for *BIN1* isoform 6, isoform 9, and isoform 10&12, respectively. * indicates *p* < 0.05.

**TABLE 1.** Demographics of patients in a Phase 2 AD trial (NCT02880956)

|  | **Placebo** | **Tila-treated** |
| --- | --- | --- |
| **Number** | 105 | 315 |
| **Age** | 71.6 +/- 6.7 | 71.7 +/- 6.5 |
| **Sex (F) (%)** | 62 (59%) | 150 (47%) |
| **Race (White) (%)** | 100 (95%) | 304 (96%) |
| ***APOE*ε4+ (%)** | 72 (68%) | 222 (70%) |

Abbreviations: Tila, tilavonemab; *APOE* ε4, apolipoprotein E ε4

To further understand the impact of the genetic variants on disease, we evaluated the association of *BIN1* rs35103166 with the change in cognitive score using the clinical dementia rating (CDR)-Sum of Box scale, which is commonly used to diagnose dementia due to AD. We specifically evaluated this change in the placebo group so we could identify any changes associated with disease progression over the 2-year timeframe of the clinical trial. We found no significant changes associated with the genotype at rs35103166 at baseline or at longitudinal time points (Supplementary S5 (F)).

## 4 DISCUSSION

We have demonstrated that transcript isoforms of *BIN1*, the second-most significant genetic risk factor for AD, are distributed in a cell-type specific manner in a hiPSC model differentiated into neurons or microglia. Whereas this reflects previous findings from other models, our work is the first to demonstrate that the expression of these isoforms changes with differentiation of hiPSCs.

Recently, the US Congress put forward the Food and Drug Administration (FDA) Modernization Act 2.0 which recommended the use of alternatives to animal testing, one of which may be hiPSC [60]. The use of hiPSC has revolutionized our understanding of disease mechanisms and has proved to be a valuable tool for developing novel therapeutics. We were able to effectively demonstrate that differentiation-associated changes in isoform expression can be monitored in an iPSC model system. This provides a foundation for using hiPSC not just to characterize neurodegenerative diseases, but also to monitor expression of specific isoforms during cellular development.

Splice isoforms of *BIN1* have been implicated in the acceleration of AD, potentially through affecting tau aggregation [61], regulating β-secretase cleavage of Aβ precursor protein (APP) and Aβ generation [39], pro-inflammation [31], synaptic dysfunction, and exocytosis of synaptic vesicles [62–64]. Our work implies a role for alternate splicing as a potential mechanism by which *BIN1* variants manifest their effect. Furthermore, we have been able to find differential isoform expression in the PB from AD patients associated with one of the *BIN1* SNPs.

The impact of variants in the *BIN1* locus has been studied extensively. A recent publication found that microglia specific enhancers harbor a non-coding *BIN1* AD risk variant, the deletion of which led to microglia-specific ablation of *BIN1* expression [65]. Loss of *BIN1* has been shown to elevate the production of Aβ42 via BACE1 accumulation at early endosomes [66]. The *BIN1* variant rs744373 has been associated with increased Tau pathology and declining memory performance in the ADNI cohort [52]. Previous work has also shown that AD brains have an overall decrease in *BIN1* neuronal isoform 1 and an increase in the microglial isoform 9 [41,42], which may be due to a shift in brain cell populations, driven by neuronal death and gliosis [31]. Our work provides support for a hiPSC model to help disentangle the changes driven by each of the cell types.

Recent publications have shown that dysregulation of mRNA splicing may be a mechanism by which the genetic risk variants mediate disease etiology. Lopes et al demonstrated that known AD risk variants in *CD33* and *MS4A6A* may act through modulation of splicing in the microglia [65]. A cross-ancestry atlas of the developing human brain demonstrated that isoform-level regulation mediated a large proportion of GWAS heritability during neurodevelopment [67]. Exploring this cross-population resource of gene, isoform, and splicing regulation demonstrated that rs35103166, the SNP associated with differential *BIN1* isoform expression in PB, was a splice-quantitative trait locus. Furthermore, this SNP is within a distal enhancer-like signature as identified by the ENCODE project [68]. Lopes et al also demonstrated that candidate causal variants for AD, e.g. USP6NL are within microglia-specific enhancers, and mediate the expression of the gene in a cell-type specific manner [65].

If the alternate splicing of *BIN1* is the primary mechanism by which *BIN1* risk variants promote disease pathogenesis, this may be a novel target for the development of drugs targeting neurodegenerative diseases [69]. Modulation of RNA splicing can be achieved by targeting the components of the spliceosome, which would impact the efficiency of splicing. Alternatively, the aberrant isoforms may be targeted, which may offer more specificity [70].

We recognize that there are several limitations to this study. We used ddPCR, a quantitative readout for the *BIN1* isoforms but it was very limited in the range of detection of splice forms. A long-read method [71], for e.g., nanopore sequencing, would provide an unbiased overview of the various splice isoforms found in the locus. A second limitation was that only PB samples were available from the clinical trial, and we were therefore only able to detect microglia-specific *BIN1* isoforms, given the low expression of neuronal isoforms in the periphery. Accessing a sample set which includes matched PB and CNS tissue would provide an understanding of the association of *BIN1* risk variants with the expression of the various isoforms. Finally, since we had a smaller number of individual samples analyzed for the clinical trial, we didn’t observe any significant associations of cognitive scores with *BIN1* genotype or isoform expression. Having a larger number of samples may power the study to detect associations between cognition and the *BIN1* genotype and the expression of *BIN1* isoforms.

## RESEARCH IN CONTEXT

1. **Systematic review:** The authors did a literature search in databases (e.g., PubMed) and reviewed the most related literature to evaluate the role of *BIN1* on AD risk, but the precise mechanisms by which *BIN1* genetic variations drive the expression of *BIN1* isoforms and its association with AD risk is still largely unknown.
2. **Interpretation:** The present study shows *BIN1* variants are significantly associated with AD risk in GWAX analysis from UKBB. Our work is the first to show that the expression of these isoforms changes with differentiation stage within the hiPSC model. We also demonstrated the influence of genetic *BIN1* risk alleles on *BIN1* isoform expression in PB from a Phase 2 AD trial (NCT02880956).
3. **Future directions:** Large AD-related cohort / sample numbers should be evaluated in the future in order to evaluate promising genes for drug development and stratification approaches.

## ACKNOWLEDGMENTS

The authors would like to acknowledge the following AbbVie employees Mengzhen Liu, Justin E Ideozu, Elizabeth Asque, Britney Milkovich and Former AbbVie employees Ameya Kulkarni and Marie-Theres Weil for their contributions to this paper.

## CONSENT STATEMENT

All authors were employees of AbbVie when the work was being conducted. Sahar Esmaeeli is currently at Neuron 23 (South San Francisco, CA) and Lindsay Stolzenburg is currently at Northwestern University (Evanston, IL). The design, study conduct, and financial support for this research were provided by AbbVie. AbbVie participated in the interpretation of data, review, and approval of the publication. No honoraria or payments were made for authorship.

## Abbreviations

GWAS: Genome-wide association study
AD: Alzheimer’s disease
BIN1: *Bridging integrator 1*
SAIGE: Scalable and Accurate Implementation of Generalized Mixed Model
hiPSC: Human induced pluripotent stem cells
SNP: Single nucleotide polymorphism
NPC: Neural progenitor cells
PB: Peripheral blood

## SUPPORTING INFORMATION

Additional supporting information can be found online in the Supporting Information section at the end of this article.

