## Supplementary Material for "New Insights on Bridging Integrator 1 Protein Isoforms as a Risk Increasing Gene in Alzheimer’s Disease"

### Supplementary Table S1

| Expressed in Cell T | Name | Forward Primer | Reverse Primer | TagMan Probe |
|---|---|---|---|---|
| Neuron[26,28] | *BIN1* isoform 1 | CCAAGGAAGTCAAGCAGGAG | AAAGTCCAGGTCCAGCAGAC | AGGACACGTTTGTCCCTGAG |
| Neuron[26,28], Astrocyte[28] | *BIN1* isoform 2 | AGAGTCAACCACGAGCCAGA | AAAGTCCAGGTCCAGCAGAC | AGTCCCCATCTCAGTTTGAG |
| Neuron[26, 28] | *BIN1* isoform 3 | GAGTCAACCACGAGCCAGAG | CCAGGACACAGCAAAGGTG | TCTCAGCCCACAGAGAGTCC |
| Neuron[26,28], Astrocyte[28] | *BIN1* isoform 5 | CCAAGGAAGTCAAGCAGGAG | CCAGGACACAGCAAAGGTG | AGGACACGTTTGTCCCTGAG |
| Neuron[26], Astrocyte[28],Microglia[26,43] | *BIN1* isoform 6 | AGGACACGTTTGTCCCTGAG | GGAGGCTGCTTCACTTGC | AGCAGAGGCCTCGGAGGT |
| Neuron[26,28] | *BIN1* isoform 7 | GAGTCAACCACGAGCCAGAG | CCAGGACACAGCAAAGGTG | TCTCAGCCCACAGAGAGTCC |
| Astrocyte[28],Microglia[26,43] | *BIN1* isoform 9 | AGAGTCAACCACGAGCCAGA | GGAGGCTGCTTCACTTGC | AAGTCCCCATCTCAGCCAG |
| Astrocyte[28],Microglia[26,43] | *BIN1* isoform 10 & 12 | GAGTCAACCACGAGCCAGAG | ATTCACAGTTGCTGGGAAGG | AGAGCTCTCTTCCTGCTGTC |

**Supplementary Figure S1**

**(A)**

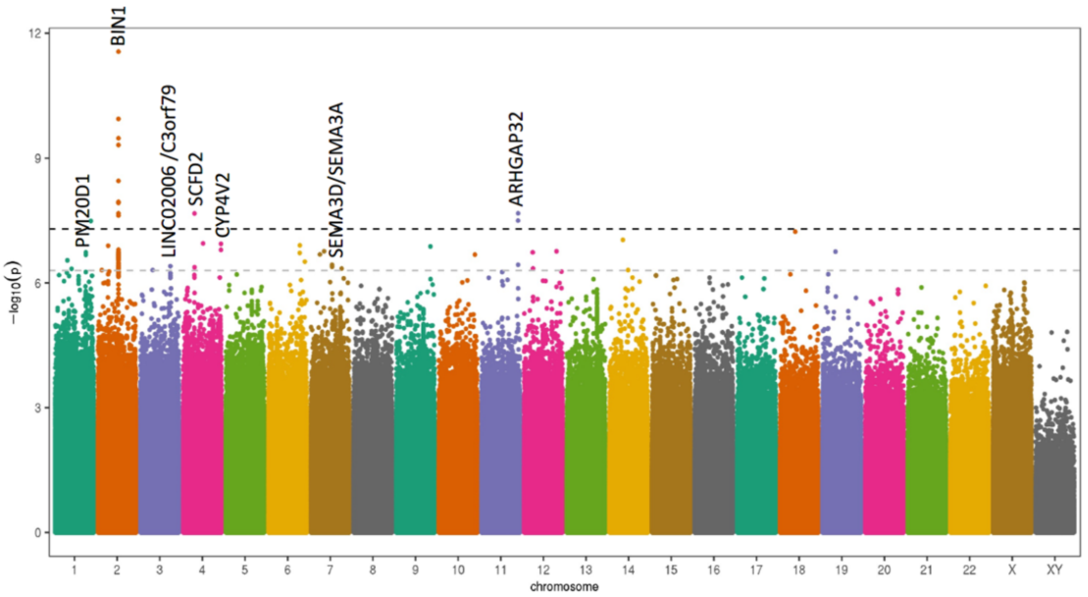


**(B)**

| | | APOE4+ | | | |
|---|---|---|---|---|---|
| **Unique ID (hg37)** | **SNP** | **pval** | **Beta** | **MAF** | **Gene** |
| 2:127892810:T:C | rs6733839 | 2.79E-12 | -0.1040716 | 0.39458055 | BIN1 |
| 11:129214412:A:G | rs560908747 | 2.11E-08 | -3.5967147 | 0.00010521 | ARHGAP32 |
| 4:53960000:G:T | rs571695314 | 2.12E-08 | -2.107774 | 0.00053917 | SCFD2 |
| 4:188541448:G:T | rs560120139 | 1.14E-07 | -2.4782731 | 0.00053131 | CYP4V2 |
| 6:139964185:G:A | rs549317591 | 1.24E-07 | -3.2072627 | 0.00020876 | TXLNB |
| 9:129461828:G:A | rs749654121 | 1.32E-07 | -3.1082817 | 0.00024491 | RALGPS1/LMX1B |
| 1:205878592:GA:G | rs538504793 | 1.84E-07 | -0.1395446 | 0.07974837 | PM20D1 |
| 7:84488029:A:C | rs192708784 | 3.67E-07 | -3.680651 | 0.00015954 | SEMA3D/SEMA3A |
| 3:153283580:C:T | rs144523018 | 3.93E-07 | -0.6815993 | 0.00311122 | LINC02006 /C3orf79 |

**Supplementary Figure S1. Stratified case-control association mapping of Alzheimer's disease using family history from UK Biobank (UKBB).** A genome-wide association study by proxy (GWAX) using imputed genotypes on SAIGE (Scalable and Accurate Implementation of Generalized mixed model) platform was conducted as described in Figure 1. In addition, data from the cases and controls were stratified based on their *APOE* status, and shown is the association study on carriers of *APOEe4*. (A) Manhattan plot with chromosome numbers on the X-axis and genome-wide significance level (-$\log_{10}$(p-value)) on the Y-axis shows most significant variants on chr. 2 at the *BIN1* locus. (B) Top significantly associated variants in an *APOE4+* stratified GWAX analysis. SNP: single nucleotide polymorphism MAF: minor allele frequency.

**Supplementary Figure S2**

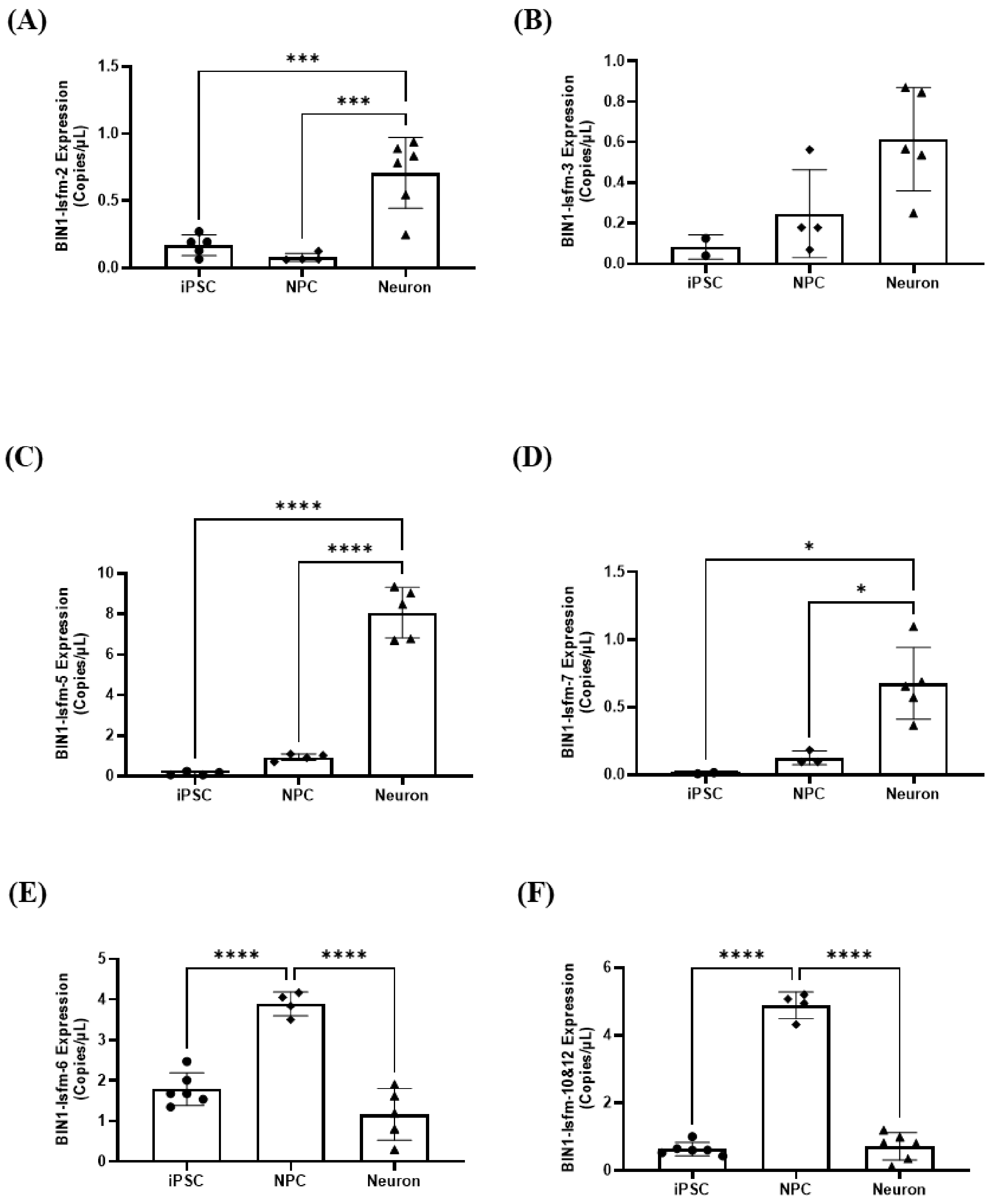

**Supplementary Figure S2**. **Change in expression level of *BIN1* isoforms during hiPSC differentiation into neurons**. hiPSCs were differentiated into neural progenitor cells (NPC), and then into neurons. The expression level of *BIN1* isoforms was measured by ddPCR technology. (A-D) Expression level of neuronal-astrocytic-specific *BIN1* isoform 2, 3, 5 and 7, respectively, during differentiation from hiPSCs into NPCs, and finally into neurons. (E-F) Expression level of microglia-specific *BIN1* isoform 6 and 10&12, respectively, during differentiation from hiPSCs into NPCs and finally into neurons. In each panel, X-axis indicates stage of differentiation, Y-axis indicates *BIN1* isoform expression quantified as copies/uL. Each point represents the mean ± S.D. of 4-6 independent experiments. Ordinary one-way ANOVA was performed first to compare between groups with ****, $p<0.0001$. Post-testing was done using Tukey's multiple comparisons test to compare every group to the other with *, $p < 0.05$; **, $p < 0.01$; ***, $p < 0.001$;****, $p < 0.001$.

**Supplementary Figure S3**

**(A)**

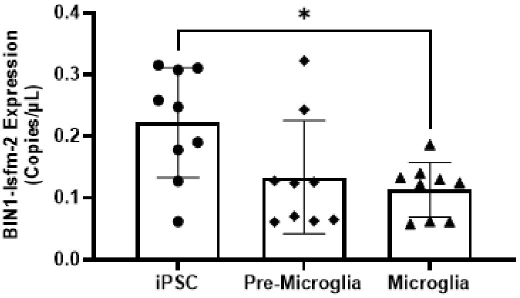


**(B)**

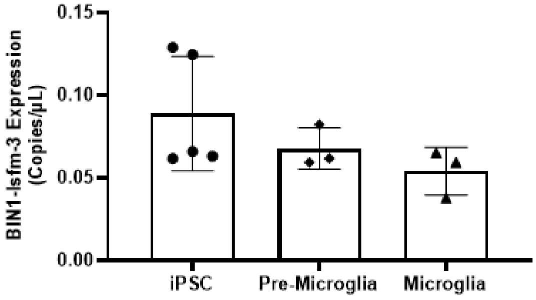


**(C)**

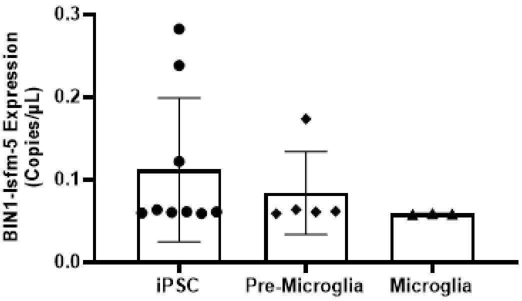


**(D)**

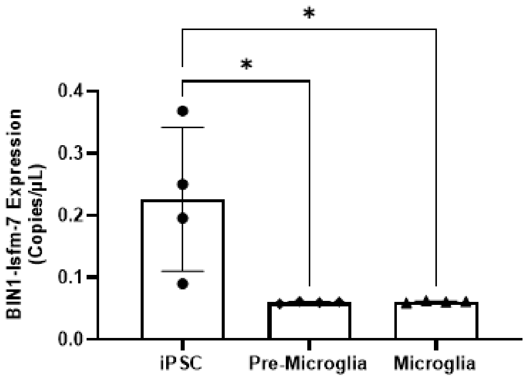


**(E)**

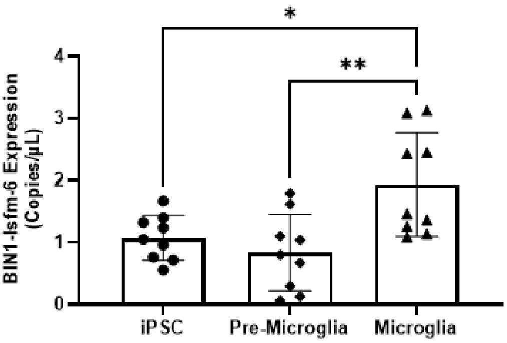


**(F)**

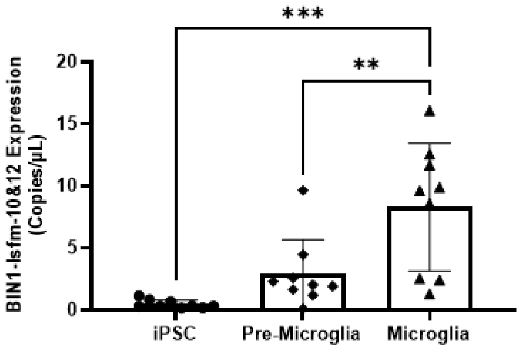

**Supplementary Figure S3**. **Changes of expression level of *BIN1* isoforms during hiPSC differentiated into microglia.** hiPSCs were differentiated into pre-microglia first, then continued differentiated into microglia. ddPCR technology was applied to detect the expression level of *BIN1* isoforms. (A-D) Expression level of neuronal-astrocytic-specific *BIN1* isoform 2, 3, 5 and 7, respectively, during differentiation from hiPSCs into pre-microglia, and finally into microglia. (E-F) Expression level of microglia-specific *BIN1* isoform 6 and 10&12, respectively, during differentiation from hiPSCs into pre-microglia, and finally into microglia. In each panel, X-axis indicates stage of differentiation, Y-axis indicates *BIN1* isoform expression quantified as copies/uL. Each point represents the mean ± S.D. of 3-9 independent experiments. Ordinary one-way ANOVA was performed first to compare between groups with ****, $p<0.0001$. Post-testing was done using Tukey's multiple comparisons test to compare every group to the other with *, $p < 0.05$; **, $p < 0.01$; ***, $p < 0.001$;****, $p < 0.001$.

**Supplementary Figure S4**

(A) (B) (C)

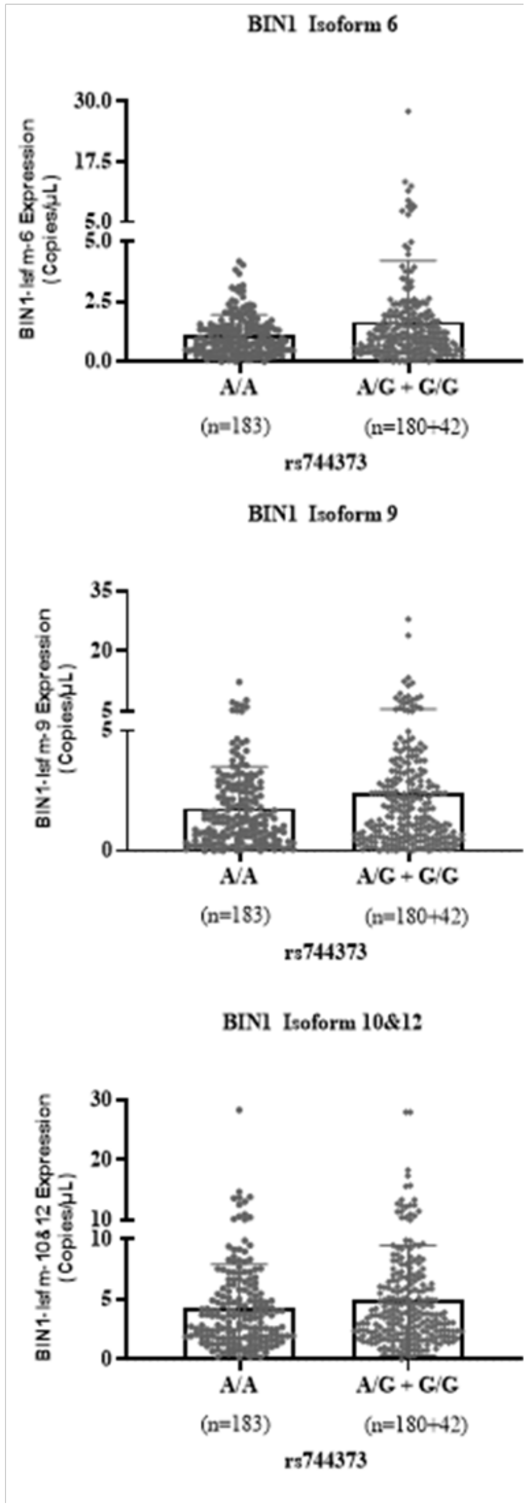

**Supplementary Figure S4. Effect of *BIN1* genotype on expression of *BIN1* isoform in peripheral blood from AD trial.** RNAs from peripheral blood samples collected at baseline from the AD trial were used to measure the expression of *BIN1* isoforms using ddPCR technology. Whole-genome sequencing data from this trial was used to identify genotype status at *BIN1* SNP rs744373. At this locus, 183 subjects had reference homozygous (A/A), 180 subjects had heterozygous (A/G) and 42 subjects had alternate homozygous (G/G) genotype. All subjects containing alternate genotypes were grouped together for this analysis, as indicated on the X- axis. Y-axis indicates *BIN1* isoform expression quantified as copies/uL. Linear regression analysis was performed to test the association of the presence of the SNP with the expression of specific *BIN1* isoforms using age, sex, race, and *APOE* status as covariates. Each individual panel shows the expression of *BIN1* isoform 6 (A), isoform 9 (B), and isoform 10&12 (C). * indicates $p<0.05$

**Supplementary Figure S5**

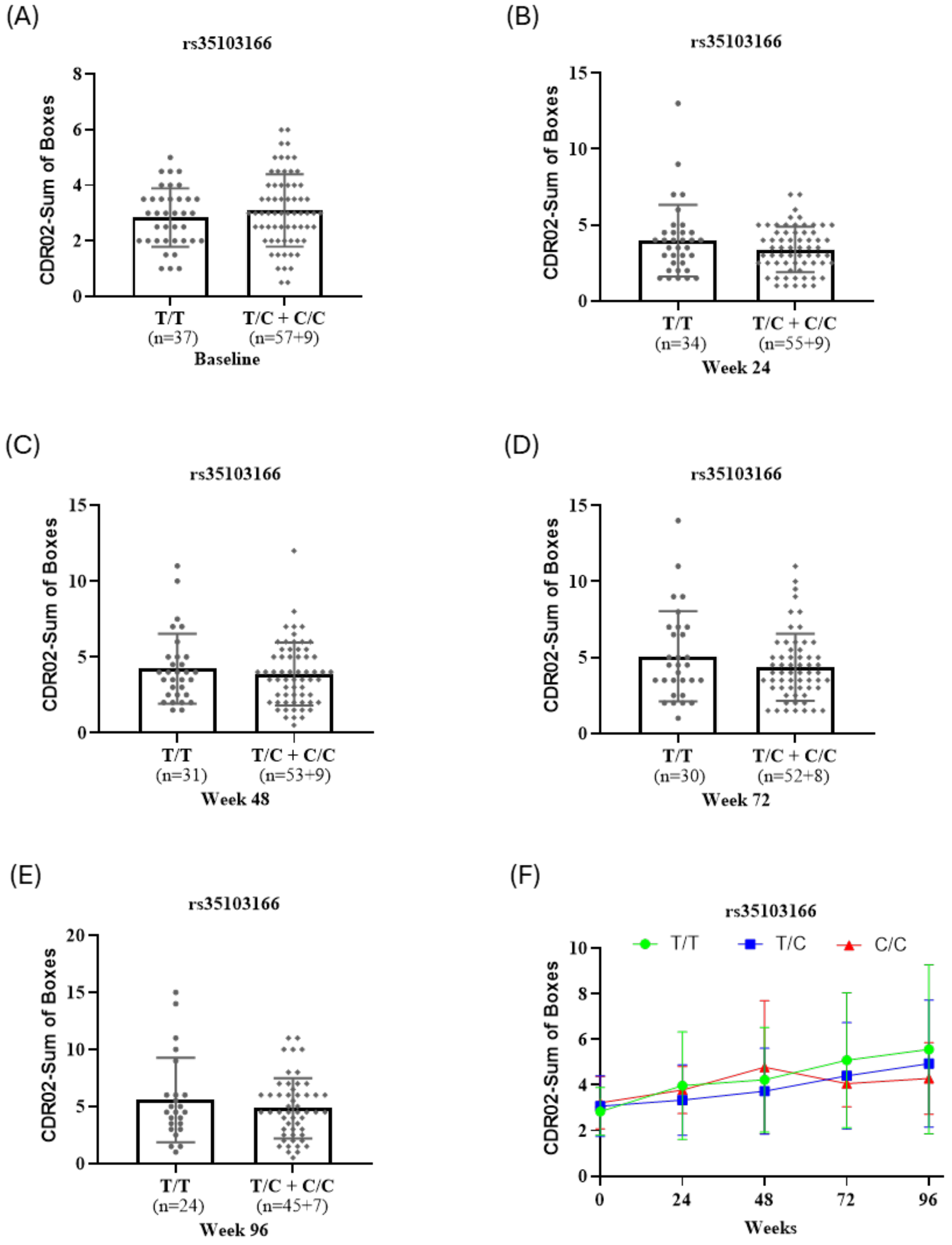

**Supplementary Figure S5 Correlation of *BIN1* genotype and AD clinical readout in PB samples from AD trial.** PB samples from placebo treated subjects were selected, and ddPCR technology was applied to evaluate the correlation of the *BIN1* SNP rs35103166 with the cognitive test Clinical Dementia Rating (CDR)02- Sum of Boxes. All subjects containing alternate genotypes were grouped together for this analysis. X-axis indicates the different visit time points. Y-axis indicates CDR02-Sum of Boxes. Linear regression analysis was performed to test the association of the presence of the *BIN1* SNP with CDR-sum of boxes at (A) baseline, (B) Week 24, (C) Week 48, (D) Week 72, (E) Week 96 and (F) across all time points using age, sex, race and *APOE* status as covariates.

### Materials and Methods

##### Human iPSC-derived neurons

Materials

| **Product Name** | **Supplier** | **Catalog Number** |
|---|---|---|
| Matrigel® Growth Factor Reduced (GFR) Basement Membrane Matrix, *LDEV-Free | Corning | 354230 |
| DMEM/F12, HEPES | Thermo Fisher | 11330032 |
| StemFlex™ Complete Medium (basal + supplement) | Thermo Fisher | A3349401 |
| RevitaCell™ Supplement (100X) | Thermo Fisher | A2644501 |
| DPBS, no calcium, no magnesium (CMF-DPBS) | Thermo Fisher | 14190136 |
| Accutase | Thermo Fisher | A1110501 |
| Essential 6 | Thermo Fisher | A1516401 |
| LDN193189 | Stemgent | 04-0074 |
| SB431542 | Tocris/R&D | 1614 |
| InSolution Cyclopamine | Millipore | 239806 |
| CHIR99021 | R&D | 4423 |

| | | |
|---|---|---|
| Advanced DMEM/F12 | Thermo Fisher | 12634010 |
| Neurobasal medium | Thermo Fisher | 21103049 |
| Neurobasal Plus medium | Thermo Fisher | A3582901 |
| Pen/Strep | Thermo Fisher | 15140122 |
| N-2 Supplement | Thermo Fisher | 17502048 |
| B-27 Supplement without vitamin A | Thermo Fisher | 12587010 |
| B-27 Plus Supplement (with vitamin A) | Thermo Fisher | A3582801 |
| GlutaMAX | Thermo Fisher | 35050061 |
| Y-27632 | Millipore | 688000 |
| Distilled water (cell culture grade, sterile) | Thermo Fisher | 15230147 |
| FGF2 | R&D | 233-FB-025 |
| DMSO | Sigma | D2650 |
| DPBS | Thermo Fisher | 14040133 |
| Laminin | Sigma | L2020 |
| RO4929097 | VWR | 10192-018 |
| Mitomycin C | Sigma | M4287 |
| BDNF | VWR | 10781-158 |
| GDNF | R&D | 212-GD-010/CF |
| Ascorbic acid | Sigma | A4544 |
| Dibutyryl cAMP (dcAMP) | Sigma | D0627 |

1) Human induced pluripotent cell lines:

In this study we used BIONi010-C-13 derived from fibroblast cells using episomal reprogramming (available through EBiSC: www.ebisc.org).

2) Manual culturing of hiPSCs with daily media change

HiPSCs were cultured in StemFlex complete medium in a T175 flask coated with 77.5 ug/mL Matrigel (Matrigel® Growth Factor Reduced (GFR) Basement Membrane Matrix, *LDEV-Free). The cells were passaged every Monday and Thursday using ReLeSR, at ratios of 1:10-1:20 depending on cell confluency. The cells were cultured in 10 mL/10 $cm^2$ StemFlex complete medium.

3) Cortical neuron differentiation

1. Generation of cortical neural progenitor cells (NPC)

HiPSC were dissociated as single cells using accutase and seeded at 500,000 cells per $cm^2$ in 2.5 mL/10 $cm^2$ of StemFlex complete media supplemented with 10μM Y-27632. The next day (Day0), when the cells were more than 90% confluent, cells were washed twice with DMEM/F12 and the media was replaced with Essential6 media supplemented with 10μM SB431542, 500nM LDN193189, and 500nM CHIR99021 at 5 mL/10$cm^2$. On Day2, the media was replaced with neural induction media, Essential6 supplemented with 10μM SB431542, 500nM LDN193189, and 5μM Cyclopamine. On day 3, the media was changed to 12ml/10 $cm^2$ of 0.5x Essential6, 0.5X N2B27-1, 10μM SB431542, 500 nM LDN193189,5μM Cyclopamine, and 1μg/mL Laminin. On Day 6 of differentiation, media was changed to 8ml/10 $cm^2$ of 0.5x Essential6, 0.5X N2B27-1, 10μM SB431542, 500 nM LDN193189,5μM Cyclopamine, and 1μg/mL Laminin. On day 8, the media was replaced with the same media as day 6. On day 9, cells were washed once with CMF-DPBS and dissociated using 1ml/10$cm^2$accutase at 37$^o$C for 15 minutes. The cells were collected in DMEM/F12, centrifuged and resuspended in N2B27-1 +10 μM Y-27632. The cells were counted and replated in a new Matrigel coated dish at 1,500,000 cells per $cm^2$ in 2.5 mL/10 $cm^2$ of N2B27-1 +10 μM Y-27632. The following day on Day 10, cells were washed with DMEM/F12, and the media was replaced with N2B27-1(50:50 Advanced DMEM F-12/neurobasal media with 1:200 N2 supplement, 1:100 B27-1 supplement 1% Penicillin/streptomycin and 1x glutamax) supplemented with 10ng/mL FGF2 plus 5μM Cyclopamine. On Day 13 the old media was replaced with N2B27-1 supplemented with 10ng/mL FGF2 plus 5μM Cyclopamine. On day 15, the media was changed to N2B27-1 plus 5μM cyclopamine. On Day 17, neural progenitor cells were frozen in 1 ml N2B27-1 media with 10% DMSO at 10 million cells/vial.

2. Differentiation of NPC into cortical neurons

NPC were thawed and plated in 1 mL of NB27-1 plus 10µM Y-27632 media at 1.33 million cells per well in a 12 well plate (Day-8). The next day the media was changed to NB27-1 supplemented with 500 nM LDN193189 and 5µM Cyclopamine. On Day -5, the media was changed to NB27-1 supplemented with 500 nM LDN193189, 5µM Cyclopamine, and 500 nM RO4929097 (RO). On Day -2, media was changed to neuronal differentiation media (N2B27-1 supplemented with 10ng/mL BDNF, 5ng/mL GDNF, 200 uM ascorbic acid, and 250 uM dcAMP). On Day-2 the plates for plating neurons were coated with 0.5mL of poly-L-ornithine. The next day poly-L-ornithine was aspirated and 1mL of 28 µg/mL Matrigel diluted in cold DMEM/F12 was added. The plates were wrapped in parafilm and stored at $4^{o}$C until the next day for final neuronal plating. Neuronal plating was performed by dissociating the cells with accutase for 20 minutes at 37C. Neurons were plated on poly-L-ornithine + Matrigel plates at 0.5 million cells per well in 1.5 mL of neuronal maintenance media (Neurobasal plus media with 1:50 B27plus, 1% Penicillin/streptomycin, 10 ng/mL BDNF, 5 ng/mL GDNF, 200 µM Ascorbic acid, 250 µM dcAMP and 1 ug/mL Laminin) plus 10 µM Y-27632 and 500 nM RO. Two days after neuronal plating, the cells were treated with 0.15 µg/mL mitomycin C to kill dividing cells. After mitomycin C treatment, ½ media changed was performed on Monday, Friday and Wednesday.

4) Collecting samples for Quantitative Real–time PCR

Cells for RNA analysis were collected in lysis buffer at hiPSC stage, Day 9 replating, Day 17 NPC, DIV0 at neuronal final plating, Day 7 neurons, Day 14 neurons, Day 28, and 6-weeks neurons.

**Human iPSC-derived microglia**

1) Human induced pluripotent cell lines:

In this study we used BIONi010-C-13 derived from fibroblast cells using episomal reprogramming (available through EBiSC: www.ebisc.org).

2) Culturing of hiPSCs with daily media change

HiPSCs were cultured in mTeSR Plus media (StemCell Technologies) on 6-well plates coated with a 1:600 dilution of Matrigel (hESC-Qualified Matrix, LDEV-free, Corning), and incubated for at least one hour at 37°C. The cells were passaged every 3-4 days using EDTA, at ratios of 1:8, 1:12, or 1:15 depending on cell confluency. To support cell survival, 10µM Y27632 dihydrochloride (ROCK inhibitor, StemCell Technologies) was added to the medium for the first 24 hours following passage. Subsequently, the medium was changed daily without ROCK inhibitor. To maintain all pluripotent or differentiated HiPSC cultures, 2ml of medium per 9.6cm² of surface area or equivalent was used. The passage medium containing the ROCK inhibitor was replaced after 24 hours.

3) Microglia differentiation

Generation of microglial-like cells was based upon the method by Haenseler et al 2017 . Briefly, embryoid bodies were generated by dissociating hiPSCs into single cells utilizing Accutase in a cocktail of 50ng/ml hBMP4 (ThermoFisher), 50ng/ml VEGF (ThermoFisher), and 20ng/ml SCF (Peprotech) with 10µM ROCKi. Cells were then plated at a density of 400,000 cells/well in 2ml mTeSR plus into 24-well Aggrewell 800 (StemCell Technologies) plates according to manufacturer's suggested protocol. Embryoid bodies were then fed every 24 hours with the same media cocktail (without 10µM ROCKi) by applying two 50% media exchanges per well for an additional 4 days.

On the fourth day, the embryoid bodies from each well were filtered to remove any debris and re-suspended in 35ml factory media comprised of Ex-Vivo 15 (Lonza), Glutamax 1:100

(ThermoFisher), Penicillin/Streptomycin 1:100 (Life Technologies), 25ng/ml IL-3 (ThermoFisher), 100ng/ml M-CSF (Peprotech) and 50uM beta-2 mercaptoethanol (BME) (Life Technologies). The cell/cytokine cocktail of each well was then gently transferred to a T175 flask. After two weeks, 20ml of media was removed, and replaced with 20ml fresh factory media. Subsequently each week, 25ml was removed from each factory flask and replaced with 25ml complete factory media. After 5-6 weekly feeds, the embryoid bodies begin to produce the macrophage precursors.

The embryoid bodies produce pre-macrophage precursors for 15-18 weeks. We pooled the pre-macrophage contents of the factory flasks weekly and counted the pre-macrophages, while retaining the embryoid bodies in the flasks. The pool of harvested pre-macrophages was then plated and differentiated into microglial-like cells.

After counting, the pre-macrophages were diluted appropriately with microglia differentiation media comprised of Advanced DMEM/F12 (Life Technologies), N2 supplement 1:100 (Life Technologies), Glutamax 1:100 (ThermoFisher), Penicillin/streptomycin 1:100 (Life Technologies), 50uM 2-mercaptoethanol (2-ME) (Life Technologies), 100ng/ml IL-34 (PeproTech) and 10ng/ml GM-CSF (PeproTech). Cells were plated into 6 well tissue culture plates at 150,000 cells/$cm^2$. Subsequently a single 50% media exchange was performed every 48-72 hours. Within one to two weeks, differentiating microglial-like cells begin to exhibit typical morphological features. Microglia-like cells were collected for RNA extraction in Trizol (ThermoFisher) on day 14 of differentiation.

Quantitative Real–time PCR

RNA isolation was performed using the Zymo Direct-zol (Zymo) Miniprep RNA extraction kit, following the manufacturer's protocol.
